# CarboLogR: a Shiny/R application for statistical analysis of bacterial utilisation of carbon sources

**DOI:** 10.1101/695676

**Authors:** Kevin Vervier, Hilary P. Browne, Trevor D. Lawley

**Affiliations:** The Wellcome Trust Sanger Institute

## Abstract

**Summary:** The Biolog Phenotype Microarray (PM) and Anaerobic MicroPlates (AN) 96-well plates utilise colorimetric redox reactions to rapidly screen bacteria for the ability to utilise different carbon sources and other metabolites. Measurement of substrate utilisation as bacterial growth curves typically involves extended data normalization, outlier detection, and statistical analysis. The *CarboLogR* package streamlines this process with a Shiny application, guiding users from raw data generated from Biolog assays to growth profile comparison. We applied chemoinformatics approaches to define clusters of carbon sources, based on molecular similarities, increasing statistical power. Altogether, *CarboLogR* is a novel integrated tool providing automatic and high-level resolution for bacterial growth patterns and carbon source usage.

**Availability and Implementation:** *CarboLogR* application can be downloaded and installed from Github repository https://github.com/kevinVervier/CarboLogR. Tutorial, data, and examples can be downloaded at https://github.com/kevinVervier/CarboLogR/vignettes.

**Supplementary Information:** Supplementary data are available at *Bioinformatics* online.

## 1 Introduction

The Biolog Phenotype MicroArrays and Anaerobic MicroPlates allow efficient measurement of carbon sources and other metabolite usage in bacterial strains. The output data from these 96-well plates can be used for bacterial identification, defining growth requirements or for understanding fundamental bacterial metabolism. Data processing and normalization are key steps in the data analysis, as even technical replicates of the same strain tend to vary in the measured intensity (Vaas *et al*., 2012). Current approaches either limit the analysis to single substrate signal (Vehkala *et al*., 2015), or only provide data processing without statistical comparison between groups (Cuevas *et al*., 2017). It has been demonstrated that additional statistical power in detecting patterns can be achieved by considering *a priori* grouping of categories (Khatri *et al*., 2012). Here we present *CarboLogR*, a standalone solution for bacterial growth curve analysis providing accessible quality control, chemoinformatics and well-powered statistical resources through a graphical user interface (https://github.com/kevinVervier/CarboLogR).

## 2 Material and methods

### 2.1 Quality control and outlier detection

Bacterial growth curve is numerically estimated using standard logistic growth model, as implemented in *Growthcurver* package (Sprouffske & Wagner, 2016). Each growth signal is adjusted for instrumental noise by subtracting the ‘blank’ well (A01) signal. Goodness-of-fit is then used to determine if growth is detected for a specific bacterium-source pair. Kinetics parameters, such as growth rate, inflection time, and doubling time, are also estimated at this step. For each bacterium, available technical replicates are used to estimate the average number of detected wells and the software reports as outlier, any replicate with an abnormal number of detected wells, as estimated by +/- 1 standard deviation. Then, for the remaining replicates, wells for which growth is not confidently detected in at least half of the replicates are filtered out. It is important to note that this data processing step is not Biolog specific and can be applied to any 96-well plate reader data.

### 2.2 Chemoinformatics analysis

The 200 carbon sources found in the Biolog PM1, PM2A, and AN plates (**Supplementary Materials**) were curated, annotated, and manually assigned to broad categories (e.g., amino acid or carbohydrates). In addition, their chemical and molecular properties were downloaded from the PubChem website using *ChemmineR* package (Cao, 2008). Tanimoto distance was used to estimate similarity between the carbon sources based on their fingerprint-based similarity (Bajusz *et al*., 2015) and single hierarchical clustering was applied to determine groups of similar sources (or chemo-cluster, **Supplementary Fig 1**). Optimal number of chemo-clusters was estimated based on the maximal average silhouette width as described in (Kaufman and Rousseeuw, 1990). Group enrichment in KEGG pathways (Kanehisa, 2015) was performed using Fisher’s exact test and over-represented categories were used as chemo-cluster descriptive features. For instance, PM1 plate chemo-cluster 8 is comprises 32 carbon sources (**Supplementary Table 1**) where KEGG pathway “*Fructose and mannose metabolism*” is over-represented (Fisher’s exact test *P*=0.016).

## 3 Results

### 3.1 Implementation

CarboLogR graphical user interface is implemented using *shiny* (Chang, 2018) and therefore is straightforward to use and does not require any programming skill (**Fig 1a**). User is required to input growth profiles, as well as a metadata file for group analysis, in the “Import plate data” panel. Then, quality control metrics are automatically extracted and potential outliers are reported to the user (“Quality control” panel). Finally, a wide range of statistical analyses is accessible via point-and-click options (“Group comparison – growth” and “Group comparison – kinetic analysis” panels).

**Fig. 1.**
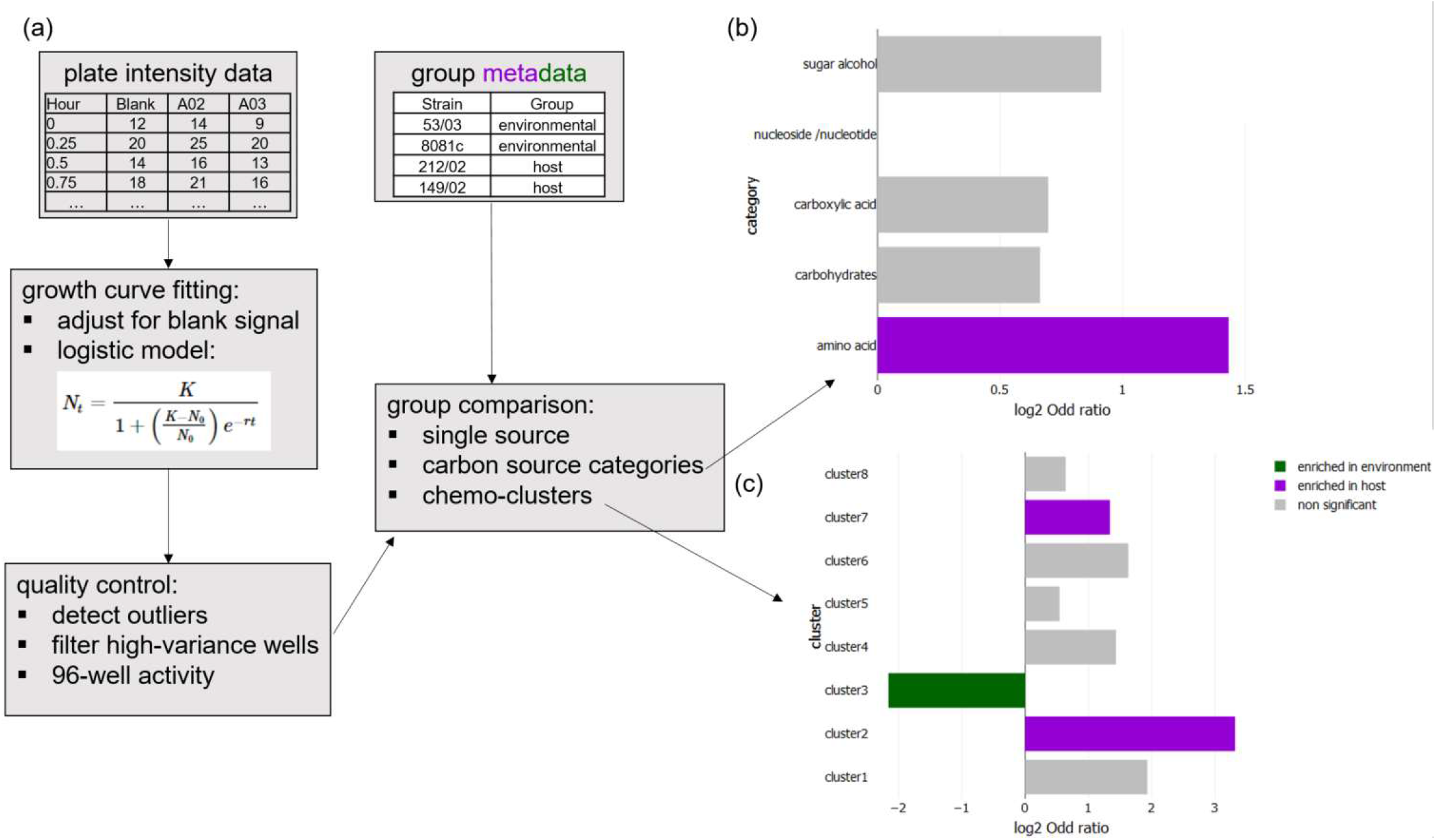
**(a)** *CarboLogR* analysis workflow: from growth profile data to statistical analyses through data quality control. **(b and c)** Group comparison of 4 *Yersinia enterocolitica* strains based on their growth profiles across 82 filtered carbon sources grouped into functional categories (b) and chemo-clusters (c). Log2-transformed odd ratio is reported as x-axis. Coloured bars correspond to significant findings (Fisher’s exact test p<0.05, green: environmental group, purple: host-specific group) and grey bars are non-significant associations.

### 3.2 Case example

First, we used data from 13 PM1 plates testing four *Yersinia enterocolitica* isolates (Reuters *et al*., 2014), two were from environmental sources (53/03 and 8081c) and two were host-specific (212/02 and 149/02). Quality control step detected and reported one outlier sample (212/02_1) as growing on 57 wells whereas the other two replicates grew on 74 sources (**Supplementary Fig 2**). The presence of growth comparison between the two strain groups did not find any significant difference at the single well level across 82 sources (**Supplementary Fig 3**), but did observe differences in the kinetic parameters. The host-specific strains grew at a faster rate (**Supplementary Fig 4**) with a shorter time to time to exponential growth (**Supplementary Fig 5**). Furthermore, set analyses done on carbon source categories (**Fig 1b**) and chemo-clusters (**Fig 1c**) highlighted significant differences between the strain groups. Categories enriched in host-specific strains involve amino acid metabolism (pathway example in chemo-cluster 2: “Alanine, aspartate and glutamate metabolism” and chemo-cluster 7: “Glycine, serine and threonine metabolism”). These functions are important for general metabolism and are expected in the host-specific strains (Neis *et al*., 2015). “Styrene degradation” (from chemo-cluster 3) is generally associated with environmental strains as styrene can be found in samples either naturally or due to industrial waste (Portune *et al*., 2015). Altogether, these analyses validate the original study and provide additional meaningful data to the user.

## Acknowledgments

We would like to thank Sandra Reuter, Theresa Feltwell, and Nicholas Thomson for sharing the *Yersinia enterocolitica* growth profiles.

## Supplementary data

**Supplementary Table 1.**
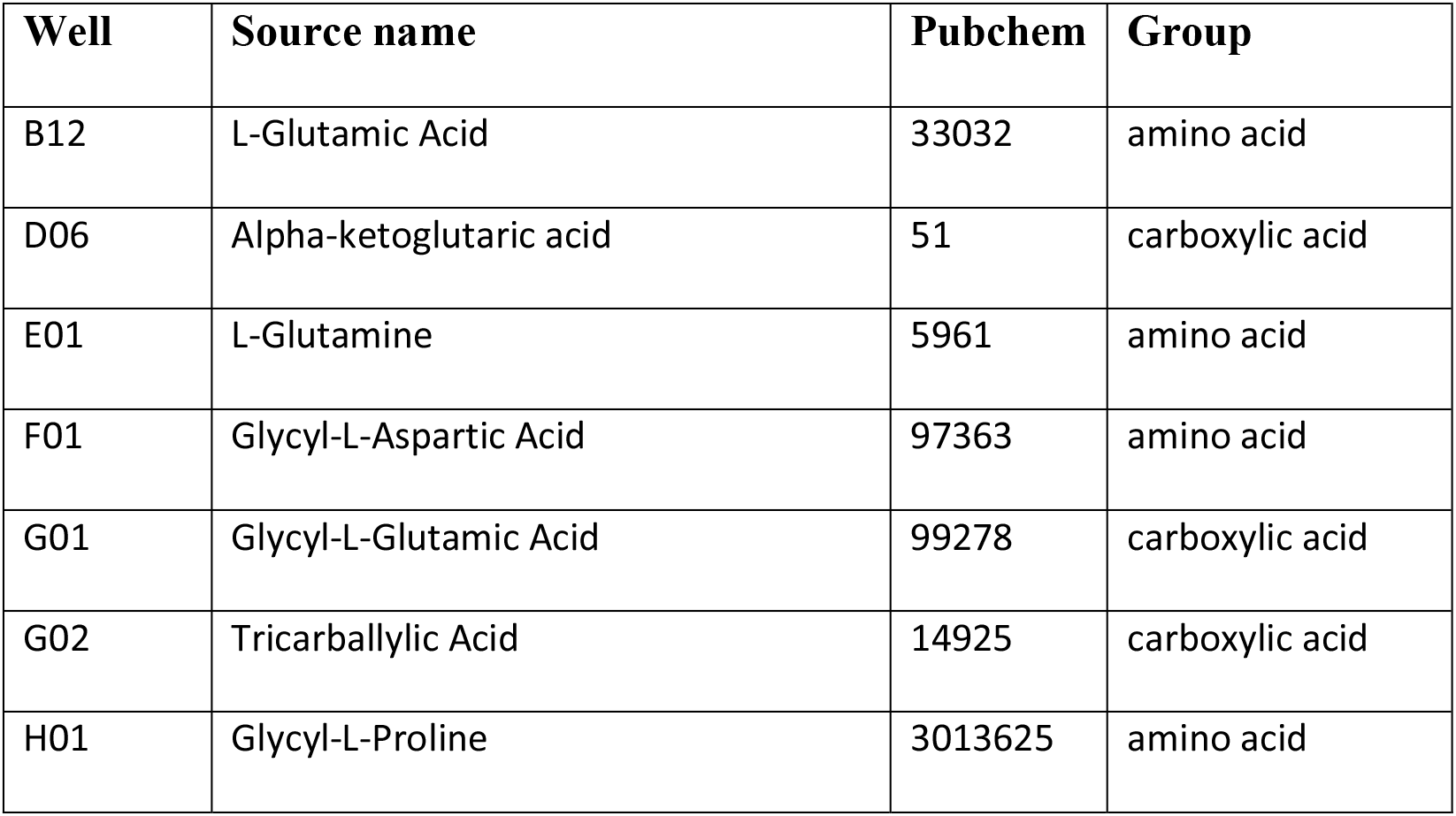
Example of chemo-cluster composition in carbon sources. PM1 plate chemo-cluster 1 comprises 7 carbon sources over-representing KEGG pathway “*D-Glutamine and D-glutamate metabolism*” (Fisher’s exact test *P*<0.01). Annotation table for all the carbon sources is available online (https://github.com/kevinVervier/CarboLogR/data/biolog_wells.txt).

**Supplementary Fig 1.**
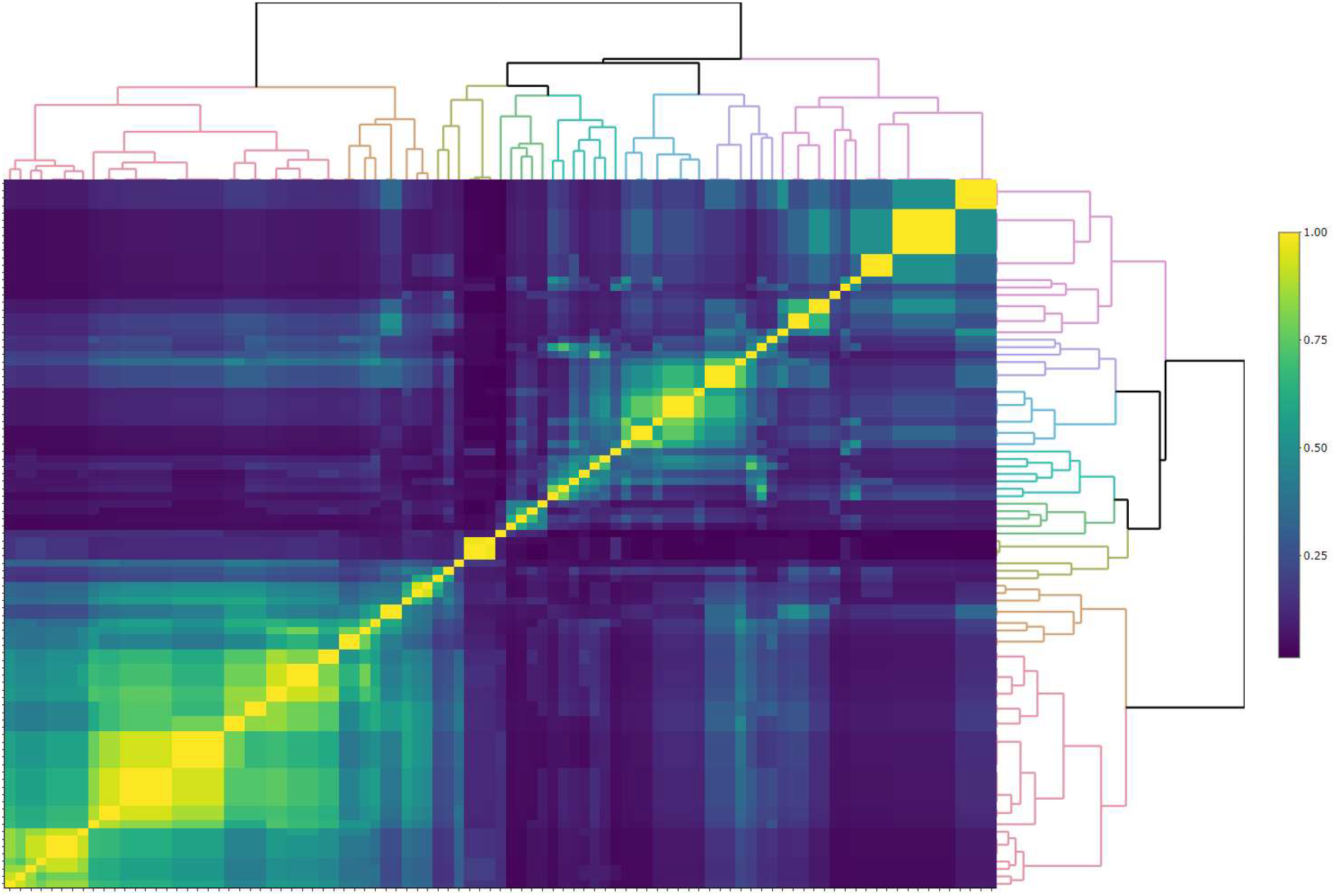
Hierarchical clustering of 96 carbon sources found on PM1 plates, based on Tanimoto distance using hashed binary chemical fingerprints as molecular representations. Cell brightness represents the degree of similarity between two sources (bright: highly similar, dark: unrelated). Each carbon source belongs to one of the 8 similarity groups (also referred as chemo-clusters), each coloured accordingly in the dendrograms. For interactive version of the heatmap, please refer to the online vignette.

**Supplementary Fig 2.**
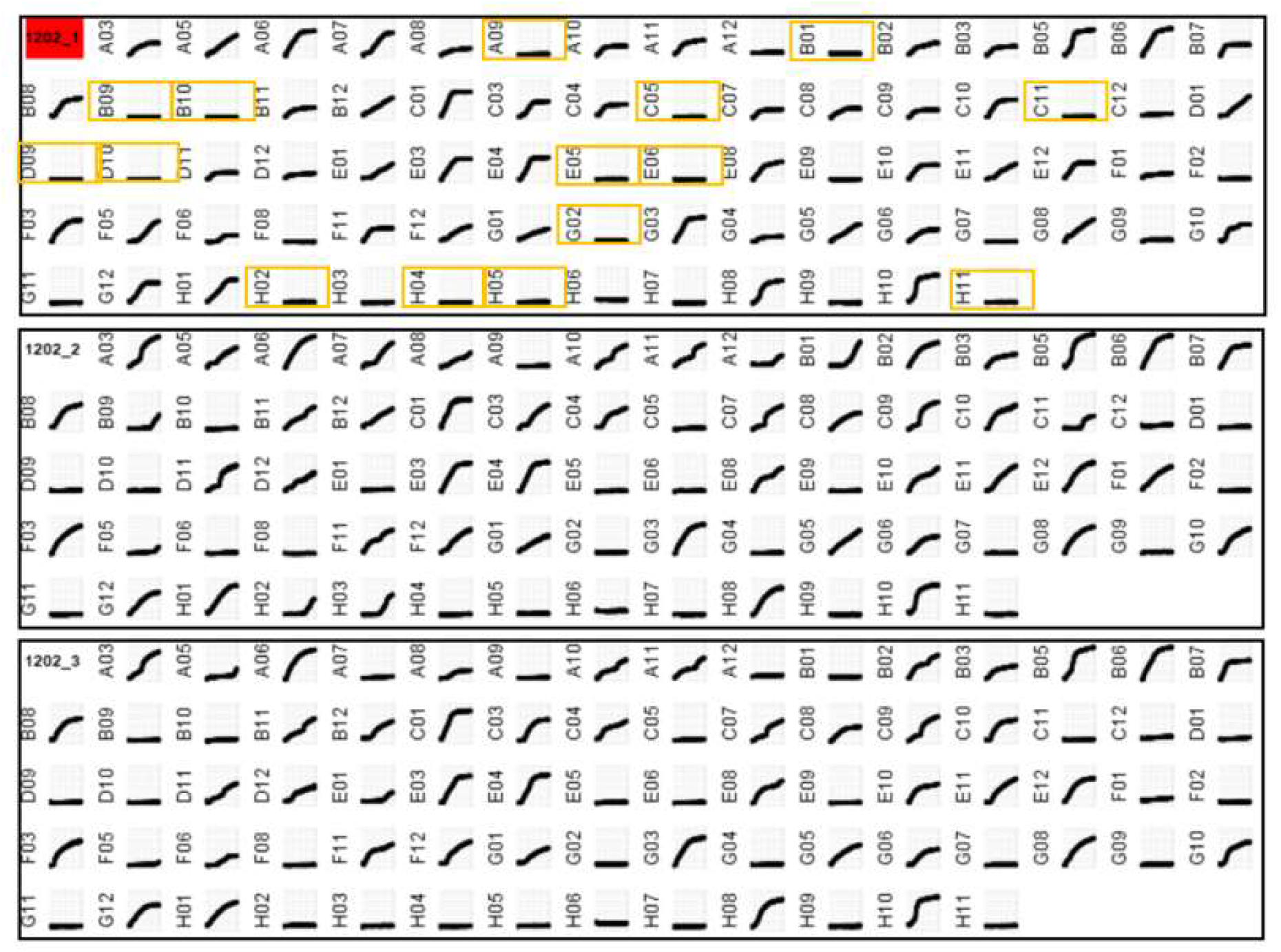
Quality control process identifies one outlier replicate (red) for strain 212/02 and displays wells stable for the majority of the replicates. Orange boxes represent carbon sources for which abnormal growth pattern was observed. Each black box corresponds to one technical replicate, and each figure represents growth pattern for one carbon source after adjusting for blank signal.

**Supplementary Fig 3.**
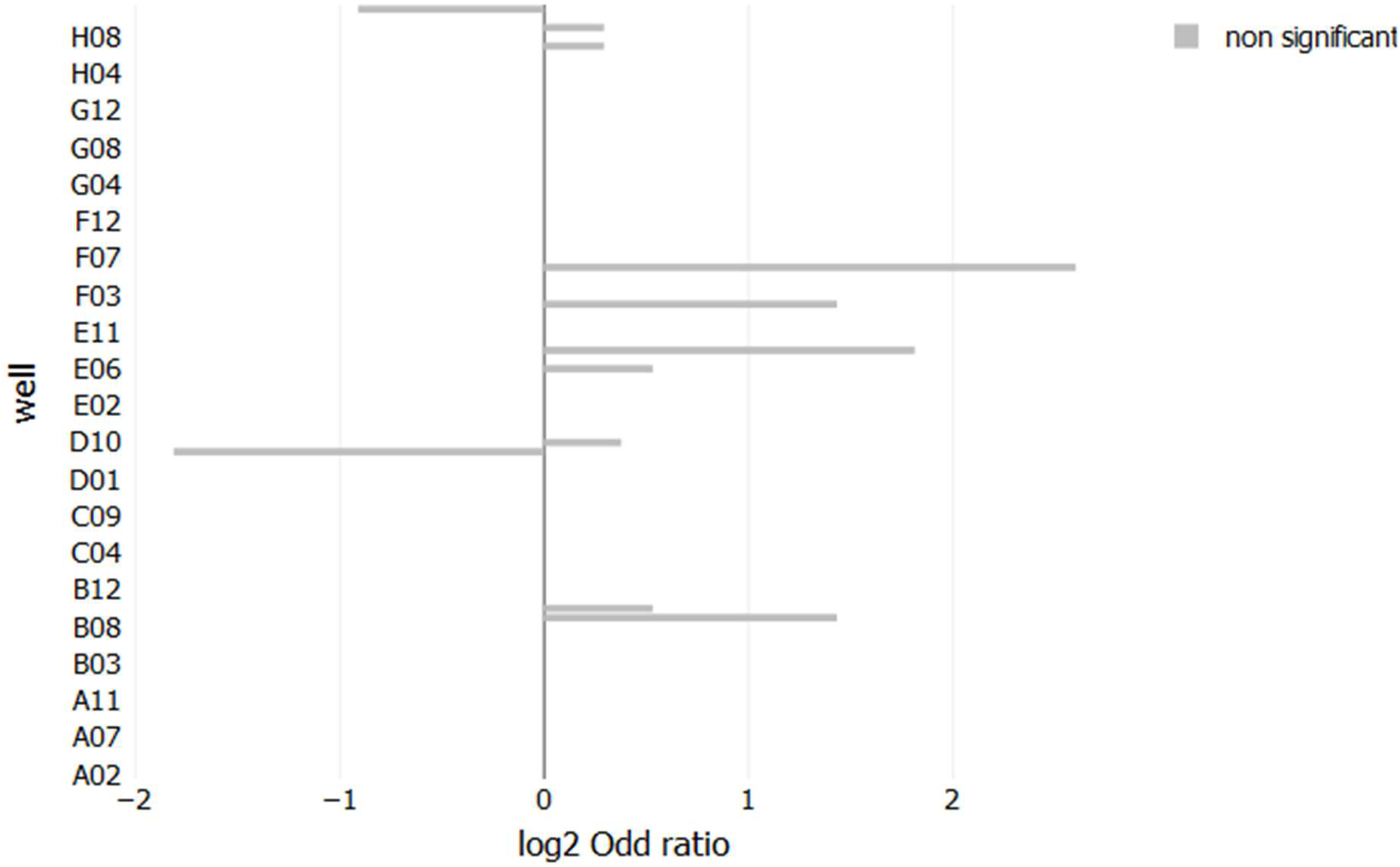
Single-well comparison of 2 *Yersinia enterocolitica* strain groups based on their growth profiles across 82 filtered carbon sources on PM1 plate. Fisher’s exact test was used to estimate how significant the difference between the two strain groups is, and log2 odd ratio is represented in x-axis. Grey bar denotes non-significant results (p>0.05). Positive odd ratio values correspond to enrichment in host-specific strains, whereas negative values relate to environmental strains.

**Supplementary Fig 4.**
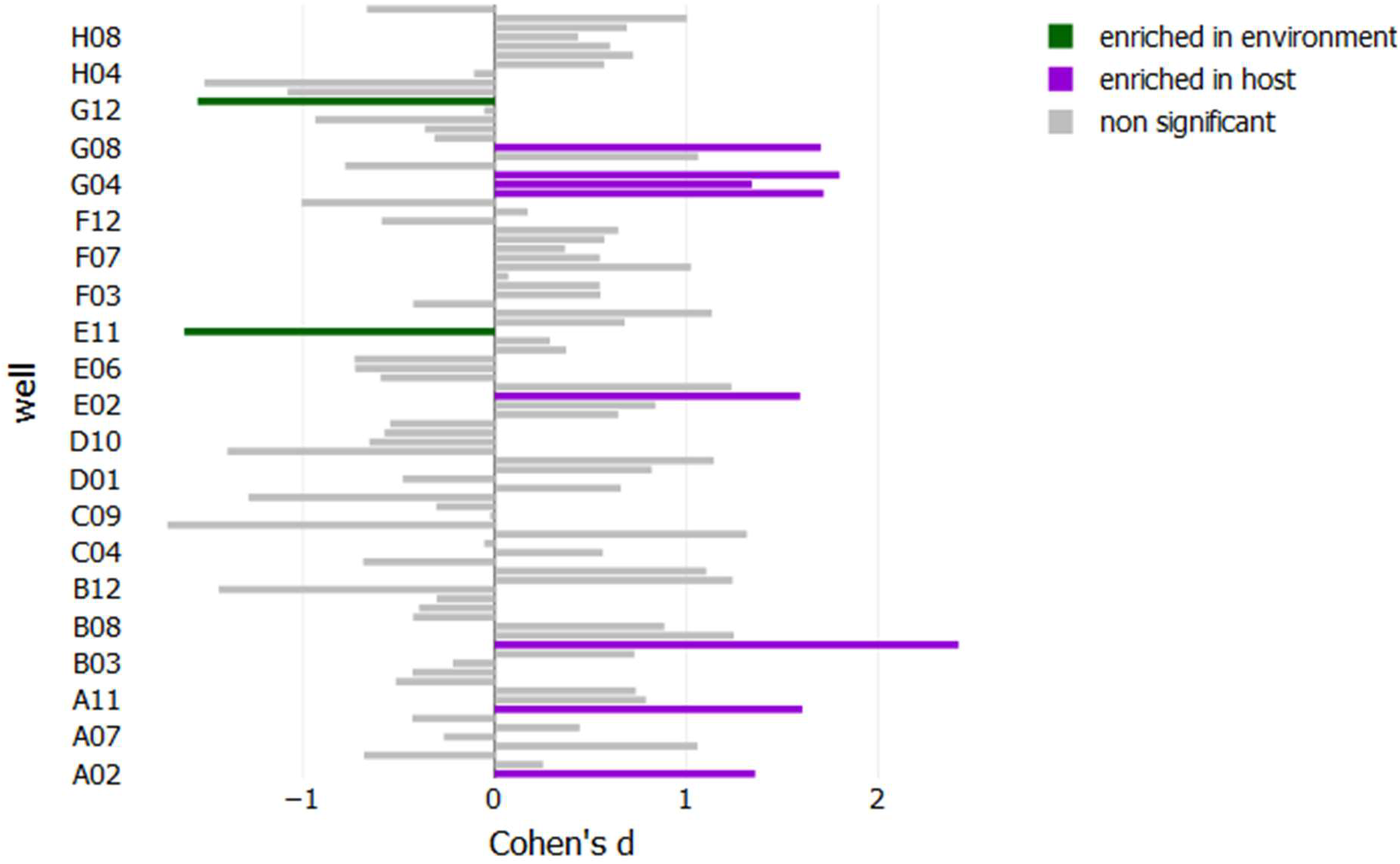
Growth rate comparison of 2 *Yersinia enterocolitica* strain groups based on their kinetic growth profiles across 82 filtered carbon sources on PM1 plate. Cohen’s *d* (x-axis) was used to estimate the effect size for each well and Student’s *t*-test was used to estimate how significant the difference between the two strain groups is. Grey bar denotes non-significant results (p>0.05). Positive effect size values correspond to faster growth in host-specific strains, whereas negative values relate to environmental strains.

**Supplementary Fig 5.**
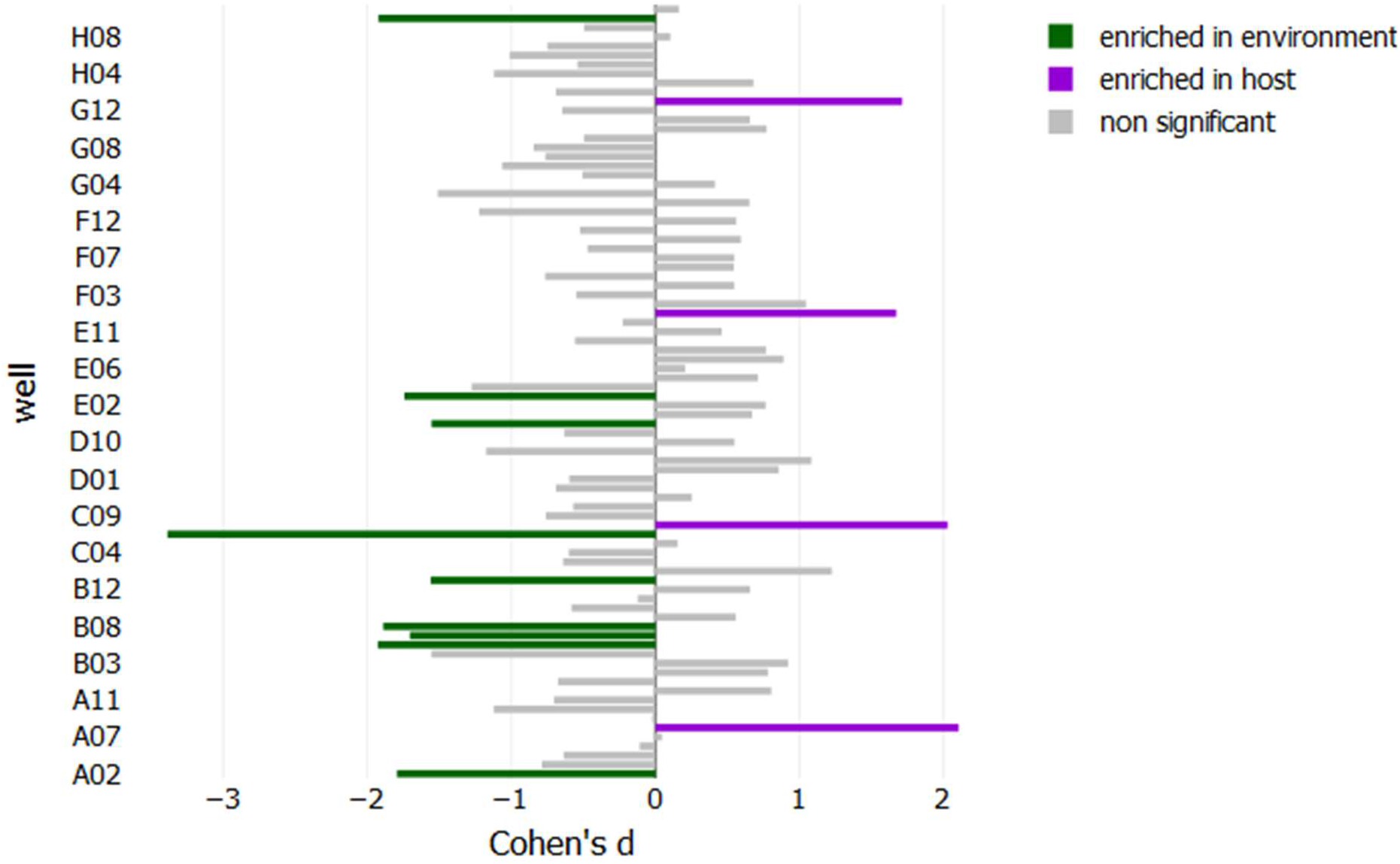
Time to exponential growth comparison of 2 *Yersinia enterocolitica* strain groups based on their growth profiles across 82 filtered carbon sources on PM1 plate. Cohen’s *d* (x-axis) was used to estimate the effect size for each well and Student’s *t*-test was used to estimate how significant the difference between the two strain groups is. Grey bar denotes non-significant results (p>0.05). Positive effect size values correspond to larger time to exponential growth in host-specific strains, whereas negative values relate to environmental strains.

